# Gibberellins target shoot-root growth, morpho-physiological and molecular pathways to induce cadmium tolerance in mung bean

**DOI:** 10.1101/2021.02.28.433230

**Authors:** Haroon Rashid Hakla, Shubham Sharma, Mohammad Urfan, Narendra Singh Yadav, Dinesh Kotwal, Sikander Pal

## Abstract

Cadmium (Cd) inhibits plant growth, perturb nutrient uptake and affect chloroplast ultra structure. Cd soil pollution is mainly contributed by excessive use of phosphate fertilizers, nickel Cd batteries, plating and sewage sludge. Research investigations deciphering role of Cd in affecting overall performance of mung bean is least understood. Likewise ameliorative effects of gibberellins (GAs) in Cd induced toxicity in mung bean are lesser known. In this context, effects of Cd stress (CdCl2, IC_50_ −500 µM L^−1^) with or without GA3 application on mung bean (*Vigna radiata* L. Var. SML-668) plants were comprehensively investigated under controlled conditions. In brief, a total of 80 mung bean plants (15 days old of uniform height) were divided into four groups, with each group (*n=20*) subjected to four different treatments (Control, CdCl_2,_ GA3, CdCl_2_+GA3), twice during the entire life cycle of mung bean plants (until harvest 85-90 days). Results revealed negative impacts of Cd stress on shoot morphometry (plant height, leaf surface area, stem diameter, shoot fresh weight, number of leaves, number of pods, length and diameter of pods), root morphometry (root length, root surface area, root dry weight, nodule number and nodule diameter), photosynthetic pigments and agronomic traits. GA3 ameliorated Cd stress by modulating shoot and root growth rates, improving overall plant metabolism, photosynthetic pigments, and shoot and root morphometry and transcript abundance of *VgPCS1*, *VgPCS2*, *VgCdR* and *VgIRT1*. Current study proposes GA3 application for the effective management of Cd induced phytotoxicity in mung bean plants.

## 1. Introduction

Heavy metal (HM) contamination of the biosphere has increased sharply for the last few years and poses major concerns for environment and human health worldwide (Hu *et al*., 2020). Metals such as cadmium (Cd), copper (Cu), lead (Pb) and chromium (Cr) could appear in natural and agricultural areas and water bodies. Latest modeling approaches showed mean bioaccumulation factors of heavy metals follow the order Cd > Zn > As > Cu > Ni > Hg > Cr > Pb in large number of crops (Hu *et al*., 2020). Among HMs, Cd pollution poses a serious threat to soil quality and human health upon consuming Cd polluted edible crops (Sanaei *et al*., 2020; Zheng *et al*., 2020). Hazardously, Cd toxicity is much higher than other organic toxic compounds due to its greater mobility (Shanying *et al*., 2017; Kubier *et al*., 2019). Cd is hardly degraded in soil, thus leads to biomagnifications in soil and plants growing in Cd polluted soils. In general, Cd is not a necessary element for plant growth and its excess has a series of harmful effects including inhibition of plant growth, perturbed nutrient uptake and chloroplast ultrastructure (Shanying *et al*., 2017; Zhao & Huang, 2018; Kubier *et al*., 2019; Shiyu *et al*., 2020). Cd generates oxidative stress in plants *via* inducing production of reactive oxygen species (ROS), such as superoxide anion radicals (O_2_), hydroxyl radicals and hydrogen peroxide (H_2_O_2_) (Azhar *et al*., 2019; Calero-Muñoz *et al*., 2019).

*Vigna radiata* L. (mung bean) is a widely grown pulse crop in Asia. It is an economical source of protein for direct human consumption (Hou *et al*., 2019). Mung bean is affected by drought (Bangar *et al*., 2019; Silambarasan *et al*., 2019), salinity (Sehrawat *et al*., 2019) and other abiotic (Ranjan *et al*., 2021) and biotic stresses (Pandey *et al*., 2018). Mung bean plants suffer significantly from HM pollution also in the agricultural soils polluted from effluents of adjoining industrial areas (Shanying *et al*., 2017; Zhao & Huang, 2018; Kubier *et al*., 2019; Shiyu et al., 2020). Reduction in plant growth and pod size occurs under HM stress in mung bean (Ghani, 2010; Wu *et al*., 2018; Shekari *et al*., 2019). Several biotechnological strategies have been adopted towards improving the HM stress management in crop plants (Ovečka & Takáč, 2014), such as HM tolerance assisted by microbes (Tiwari & Lata, 2018), transgenics (Ai *et al*., 2018; Belykh *et al*., 2019) and use of phytohormones (Sytar *et al*., 2019; Nguyen *et al*., 2021). A strong criticism for transgenic approaches of improving yield and abiotic or biotic stress tolerances of crop plants have impeded the acceptance of transgenic crops by the populace world over (Shukla *et al*., 2018; Karky & Perry, 2019). Eco-friendly approaches (phytohormonal and microbe assisted) of improving HM stress tolerances in crop plants offers an alternate way without fiddling the genetic make of crop plants and germplasm erosion (Mohite *et al*., 2017; Sytar *et al*.,2019; Haq *et al*., 2020). In mung bean seedlings, indole-3-butyric acid was shown to enhance Cd stress tolerance by improving antioxidant defense system and by promoting adventitious rooting (Li *et al*., 2018). Literature survey showed scant information about the holistic impact of Cd stress on morphometerical, reproductive, physiological and agronomical attributes on mung bean plants. Similarly, role of gibberellins (GAs), as a phytohoromonal entity for reducing Cd associated stress in mung bean plants is least explored, and mostly confined to photosynthesis (Hasan *et al*., 2020) and seedlings stages (Asgher *et al*., 2015). Current study thus comprehensively investigated the effects of Cd stress on the mung bean plants and *vis-à-vis* role of gibberellic acid (GA3-a type of gibberellins) in mitigating Cd induced stress markers. Keeping the current understanding of Cd stress in mung bean plants, present study was carried out to observe impact of Cd stress on growth as well as biochemical parameters of plants of *V. radiata* from seedling to fruiting stage in presence of 500 µM/L GA3 concentration and to provide a theoretical basis for the risk assessment of heavy metal pollution and the maintenance of sustainable agricultural production.

## 2. Material and methods

### Experiment 1: Establishment of young plants in hydroponics

Certified seeds of *Vigna radiata* L. (var. SML-668) procured from the Agriculture station, Jammu. To determine the impact of Cd stress on the establishment of root and shoot system in young plants, uniform seedlings raised for five days by paper roll method on plain water (control), were subjected to four different treatment regimes viz. Control, CdCl_2_, GA3 and CdCl2+GA3 for a period of 10 days. Young plants were harvested separately into shoot, root and emerging leaves to measure root and shoot length, and number of emerging leaves and cotyledonary leaves.

### Experiment 2: Establishment of plants in soil

Experiments were conducted in poly house (32.72°N 74.85°E) of the Department of Botany, University of Jammu, India. Certified seeds of *Vigna radiata* L. (var. SML-668) were sown in the field beds, at a uniform distance during the month of September 2018 and 2019. At 10 days after sowing (DAS), uniform healthy seedlings were selected and transplanted to pots (26 cm height and 25 cm diameter) filled with 05 kg garden soil mixed with farmyard manure in the ratio of 3:1. A total of 80 mung bean plants (15 days old of uniform height) were divided into four groups, with each group (*n=20*) subjected to four different treatments (Control, CdCl_2,_ GA3, CdCl_2_+GA3), for the next 45 days with and without treatment of GA3 (500 µM /L) by soil drench method. Water was applied through surface irrigation at the three days interval based on the calculated evapotranspiration demand following FAO-Penman Monteith model as described elsewhere (Mohan *et al*., 2019). Plants were then thoroughly investigated for changes in morphometrical, physiological parameters and molecular pathways. Prior to CdCl2 application, IC_50_ was calculated for mung bean seedlings grown in Petri dishes over a five days period.

### 2.1 Shoot and root morphometry and biomass

Number of cotyledonary and emerging leaves, seedling shoot length (SSL), seedling shoot biomass (SSB), seedling root biomass (SRB) and total root length (TRL), lateral root number (LRN) and TRL/R was measured for young plants grown in hydroponics system. Young root images were analyzed by Image J software ().

For mature plants grown in soil, shoot height (cm), stem thickness (cm), leaf number (N) and leaf surface area (cm^2^) were measured on weekly basis. Fresh and dry weights of leaves, stem and roots were measured at the time of harvest (Urfan *et al*., 2020). Root length, root surface area, nodule number and nodule diameter was was measured at harvesting stage. Root growth was imaged using a high resolution camera (Nikon Coolpix B500 Camera).

### 2.2. Histological localization and estimation of Cd accumulation by atomic absorption spectrometry

Histological localization of Cd^2+^ ions was performed with 0.5 M oxalic acid and visualized as white crystalline precipitates. Thin transverse sections of stem were dipped for 10 minutes in solution of 0.5 M Oxalic acid at a well defined ratio (?), at a controlled temperature and pH, under mechanic movement. The sections were imaged using a high resolution camera (Nikon B500) **(Supplementary Figure 1)**. For the measurement of Cd content in mature plants, atomic absorption spectrometry (AAS, Shimazdu, Japan) was used (Bakkali *et al*., 2009).

### 2.3 Biochemical estimations and stress indices

About 500 mg of fresh leaves were homogenized with ethanol (95%) (5ml) to make slurry, samples were centrifuged at 13000 rpm for 10 min and estimation of Chl *a*, Chl *b*, and carotenoids (CAR) (Lichtenthaler, 1987), sucrose content (a major photosynthate) (Miller, 1959) were carried. Protein estimation, ascorbic acid (ASA), glutathione (GSH) and total phenols (TP) were performed as described in Pal *et al*. (2016). Urease enzyme (EC 3.5.1.5) activity was measured by Berthelot method as described in Kandeler & Gerber (1988) in the root nodules. Estimation of stress indices such as proline (PL) and malondialdehyde (MDA) were performed as described in Choudhary *et al*. (2012).

### 2.4 Agronomic traits

Economically important traits of mung bean plants such as pod diameter, pod length, pod fresh weight, pod dry weight, seed number, 100 seeds weight, seeds weight/pod, seed SA, seed diameter, seed number/plant, seed protein and seed sugar content were measured at the harvesting stage using appropriate standard procedures as described elsewhere (Haider *et al*., 2020). Seed attributes particularly, width, length, thickness, surface area, Ew, Ev, Dg, Φ%, V, %G and SPr and STPC were measured.

### 2.5 Quantitative PCR analysis

Total RNA was extracted from frozen root samples using TriZol (Ambion, USA) with DNase treatment according to the manufacturer’s instruction. One microgram of RNA was reverse transcribed using ‘Transcriptor High Fidelity cDNA Synthesis Kit’ (Roche Diagnostics, Mannheim, Germany) with oligodT as a primer. Quantitative polymerase chain reaction (qPCR) analysis was conducted on Roche Light cycler®96 (Roche Diagnostics, Mannheim, Germany) and amplification curve was checked with ROCHE software version 1.1. The reaction included SYBR green PCR master mix (light Cycler 480 SYBR Green I Master, Roche Diagnostics) with 4-folds diluted cDNA in the total volume of 20*μ*l (Pal *et al*., 2016). A comparative delta Ct (2–ΔCt) method was used to quantify relative expression of genes with elongation factor (EFμ1) as an endogenous control. Primers sequences are given in (Muneer *et al*., 2014) **(Table M1)**.

### 2.6 Pearson correlation analyses

Pair-wise correlation between Cd stress with or without GA3 on plant fitness and agronomic attributes was carried out. Pearson’s correlation coefficient (*r*) was calculated using IBM SPSS 20.0 for ten traits (shoot height, stem thickness, leaf number, leaf surface area, root length) of plant fitness and agronomic importance (seed number, seed weight, seed weight/pod, seed surface area and total GSH). A linear correlation coefficient (*r*) was calculated to understand interaction of two factors at a time at *p <* 0.05 and *p <* 0.01 with number of biological replicates (*n =* 20 except for total GSH, *n* = 5) for following combination of treatments CN versus Cd, Cd versus Cd+GA3 and CN versus Cd+GA3. The *r* values fall in the range of +1 to −1, with positive values indicating positive linear relationship among two treatments and –ve value indicating the negative relationship among treatments.

### 2.7 Statistical analysis

Experiments were arranged in a randomized factorial design (2018 and 2019). Data obtained were analyzed by ANOVA and all means were separated at the *p <* 0.05 level using the Duncan test. All calculations and data analyses were performed using the IBM SPSS 20.0 for Windows software package and the values were expressed by means and standard error (SE).

## 3.0 Results

### 3.1 GA3 improves establishment of young shoot-root system under Cd stress

Establishment of young root system, cotyledonary leaves and emergence of young leaves were affected under Cd stress. Development of cotyledonary leaves is a pre-requisite for establishment of a healthy young plant. Cd stressed plants showed 13% and 74% reductions in establishment of cotyledonary leaves and emergence of young leaves compared to control plants **(Figure 1a)**. GA3 applied to Cd stressed plants showed improvement in establishment and emergence of cotyledonary leaves and young leaves by 15% and 17% respectively compared to Cd alone treated plants. GA3 alone showed 18% reduction in emergence of young leaves over control **(Figure 1a)**. Seedling shoot length (SSL) reduced by 13% in Cd stressed plants over control. While GA3 could improve SSL by 20% in Cd stressed plants compared to only Cd treated plants. About 17% increase in SSL was noted in GA3 alone treated plants compared to control **(Figure 1b)**. Seedling shoot biomass (SSB) and seedling root biomass (SRB) were reduced by 29% and 15% in Cd stressed plants over control. GA3 improved SSB and SRB by 22% and 21% respectively in Cd stressed plants compared to only Cd treated plants. No significant change in SSB and SRB occurred in GA3 alone treated plants over control plants **(Figure 1c)**.

**Figure 1.**
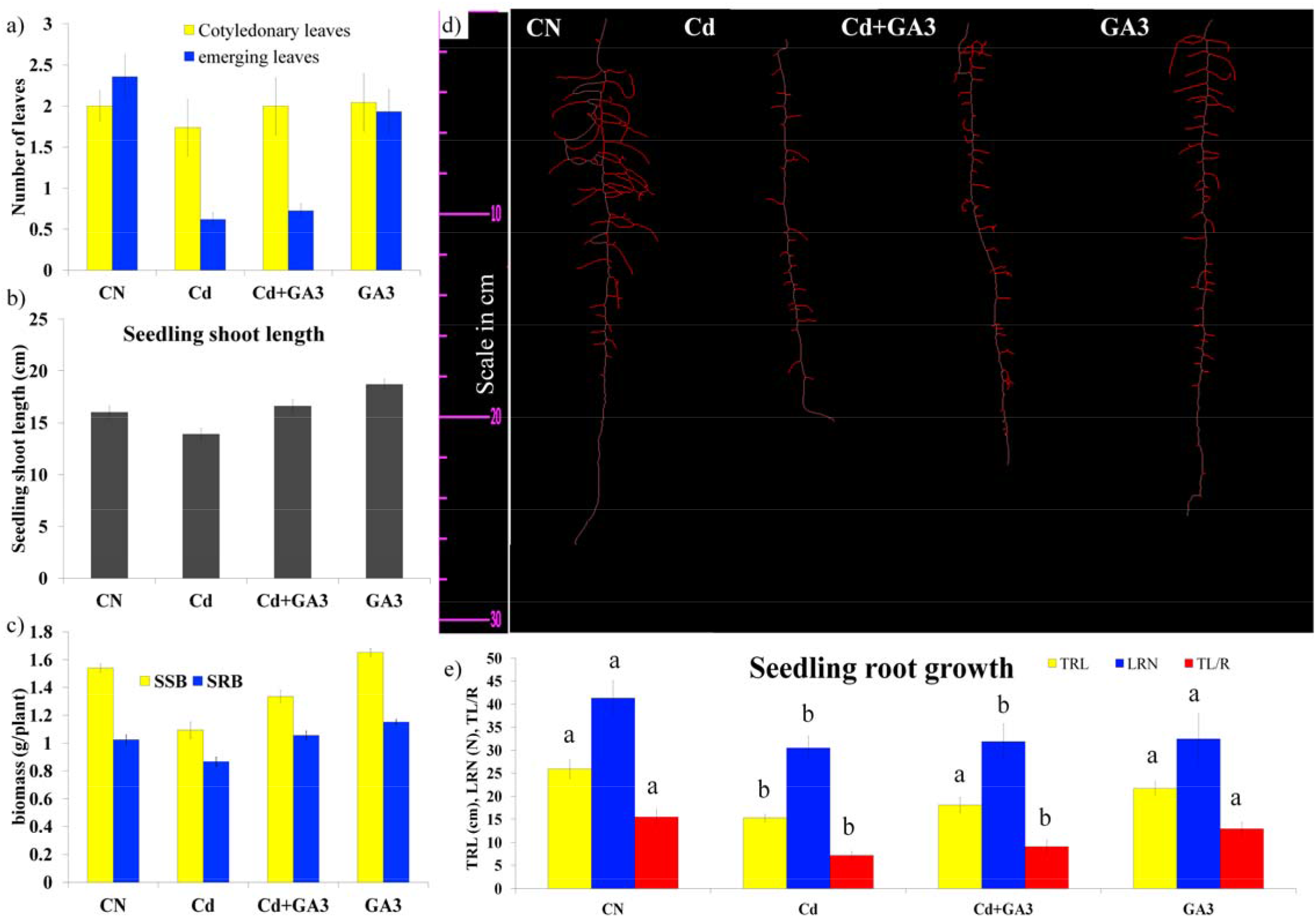
Gibberellins (GA3) improves establishment of young mung bean plants under cadmium (Cd) stress in hydroponics. Effect of GA3 with or without cadmium (Cd) stress on the number of cotyledonary and young emerging leaves (a), seedling shoot length (b), seedling shoot biomass (SSB) and seedling root biomass (SRB) (c), length of root system (d), total root length (TRL), lateral root number (LRN) and TLR/R (e) in paper roll method grown under control (CN), cadmium (Cd) stress, Cd+ gibberellins (GA3) and GA3 alone conditions hydroponically. Data presented are means ± standard errors (*n* = 10, biological replicates). Different letters (a, b) indicate significant differences from control in all combinations (Tukey’s test, *P* ≤ 0.05).

For root growth, Cd stressed plants showed significant impact on establishment of young root system, such that 41%, 26% and 54% reductions in total root length (TRL), lateral roots number (LRN) and total root length/ roots (TL/R) was recorded in Cd stressed plants over control. GA3 in Cd stressed plants showed improvements in TRL (18%) and TL/R (27%) over Cd alone treated plants. GA3 alone brought reductions in TRL, LRN and TL/R by 16%, 22% and 17% compared to control plants **(Figure 1d-e)**.

### 3.2 GA3 improves shoot morphometery under Cd stress in pot experiments

Consistent decrease in stem height and stem diameter was noted in Cd treated plants **(Fig 2a-b)**, with maximum reductions at 80 DAS and 20 DAS respectively. GA3 steadily improved stem height and stem diameter in Cd treated plants, with maximum increase recorded at 70 DAS and 20 DAS over Cd treated plants **(Fig 2a-b)**. GA3 alone improved stem height and stem diameter in a non-significant manner. A constant decrease in leaf number was noted in Cd treated plants; with maximum reduction at 80 DAS compared to controls **(Fig 2c)**. GA3 application to Cd treated plants could ameliorate negative Cd impact on leaf number, with most significant increases at 70 and 80 DAS over Cd alone treated plants. No significant change in leaf number was recorded for GA3 alone **(Fig 2c)**. At all observation points, leaf surface area (LSA) was reduced in Cd treated plants, with maximum reduction recorded at 60 DAS over control plants. GA3 improved LSA in Cd treated plants most significantly at 60 DAS over Cd treated plants **(Fig 2d)**. Increased LSA was recorded in GA3 alone treated plants significantly at 75 DAS over control plants **(Fig 2d)**.

**Figure 2.**
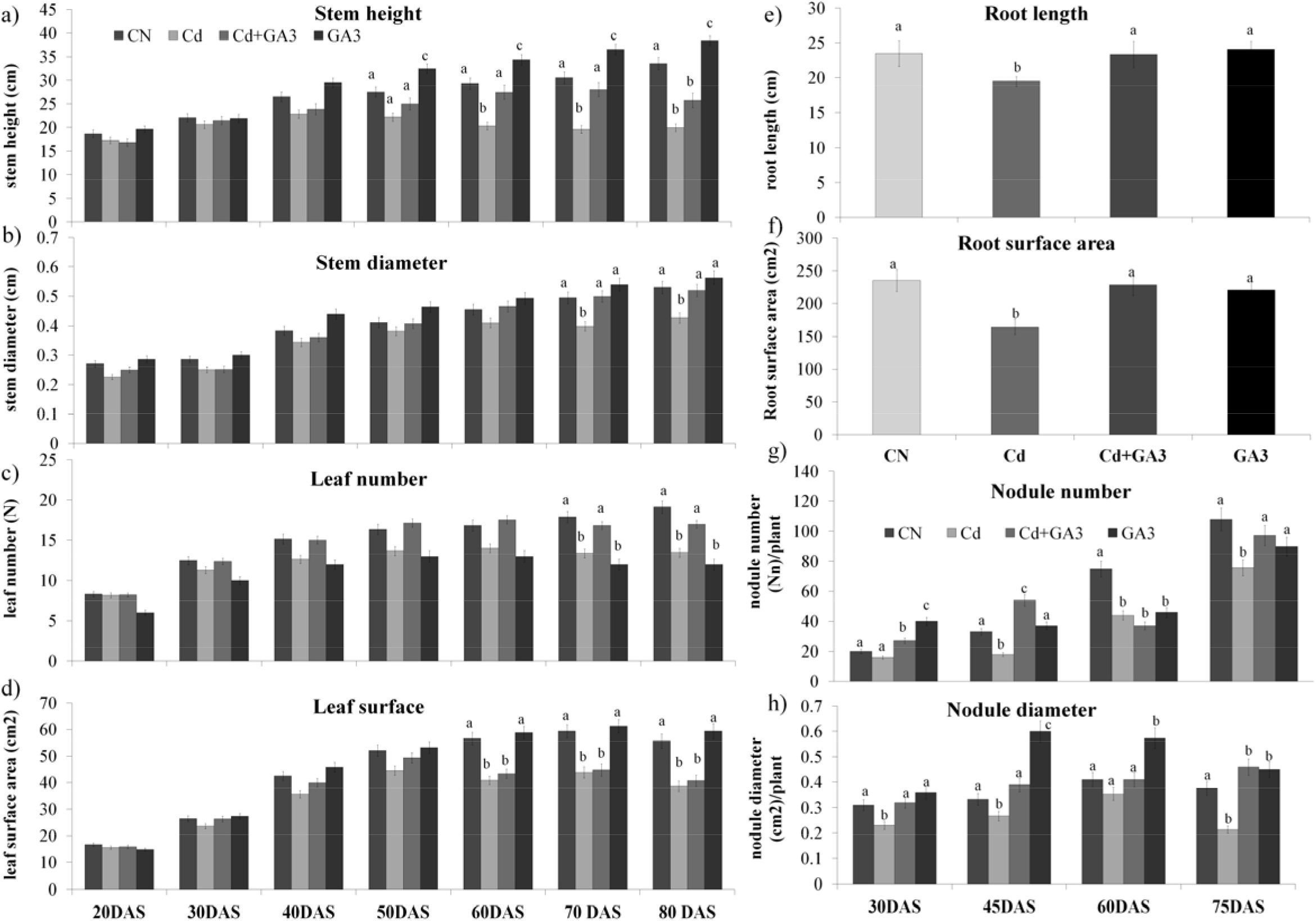
Gibberellins (GA3) alter growth parameters of mung bean plants under cadmium (Cd) stress in soil. Effect of GA3 with or without cadmium (Cd) stress on stem height (cm) (a), stem diameter (cm) (b), leaf number (*N*) (c), leaf surface area (cm^2^) (d), root length (cm), root surface area (cm^2^), nodule number (e), nodule diameter (cm) (f) of mung bean plants grown in soil in pots at 20 days after sowing (DAS), 30 DAS, 40 DAS, 50, DAS, 60 DAS, 70 DAS and 80 DAS for (a-d) and 30 DAS, 45 DAS, 60 DAS and 75 DAS for e-f grown under control (CN), cadmium (Cd) stress, Cd+ gibberellins (GA3) and GA3 alone conditions. Data presented are means ± standard errors (*n* = 10, biological replicates). Different letters (a, b, c) indicate significant differences from control in all combinations (Tukey’s test, *P* ≤ 0.05).

### 3.3 GA3 modulates root surface area and root nodules attributes under Cd stress in pot experiments

Root length (RL) and root surface area (RSA) play important role in absorption of water and uptake of minerals from the soil solution. In current study, Cd stressed plants showed 30% reduction in RSA over control plants. Furthermore, GA3 applied to Cd stressed plants reduced RSA significantly over Cd stressed plants **(Fig 2e-f)**. Increase in RSA under GA3 alone was significant compared to control plants **(Fig 2e-f)**.

Root nodule number (RNN) in Cd stressed plants increased at all the observation points (30, 45, 60 and 75 DAS) with maximum increase in RNN at 75 DAS over control. GA3 improved RNN in Cd treated plants by 35% and 64% at 30 and 45 DAS, while declined RNN by 51% and 47% at 60 and 75 DAS compared to Cd stressed plants **(Fig 2g)**. GA3 alone enhanced RNN at all observation points with maximum increase (250%) recorded at 30 DAS over control **(Fig 2g)**. Root nodule diameter (RND) showed consistent decline in Cd stressed plants, with maximum reduction (43%) noted at 75 DAS over control **(Fig 2h)**. GA3 improved RND in Cd stressed plants most significantly (46%) at 45 DAS over Cd stressed plants. RND also got improved in GA3 alone treated plants, with maximum increase (82%) observed at 45 DAS over control plants **(Fig 2h)**.

A consistent reduction in legHb content was recorded in nodules of Cd stressed plants at all observation points (30, 45, 60 and 75 DAS), with maximum decline 57% noted at 60 DAS over control. GA3 could improve legHb contents in Cd treated plants at all the observation points, with maximum increase 253% recorded for 60 DAS over Cd stressed plants **(Table 1)**. Furthermore, GA3 alone reduced legHb content at 30 and 40 DAS, while increased it by 22% and 34% increase at 60 and 75 DAS compared to control **(Table 1)**. Urease enzyme activity in root nodules decreased under Cd stress at 30, 45 and 60 DAS, while a non-significant increase in activity at 75 DAS occurred compared to control. GA3 could improve Urease activity in Cd stressed plants in a consistent manner over Cd treated plants **(Table 1)**.

**Table 1.** Effects of cadmium (Cd) stress with or without gibberellins (GA3) on the physiological attributes of leaves of *Vigna radiata* L. on chlorophyll a (Chl a), chlorophyll b (Chl b), carotenoids (CAR), total soluble sugars (TSS), malondialdehyde (MDA,), total phenol content (TPC,), total glutathione (Total GSH,), protein content (), urease activity (UA,) and nodule leg hemoglobin content (Leg Hb,) in leaf tissues of mung bean plants subjected to control (CN), cadmium stress (Cd), gibberellins (GA3), Cd+GA3 at 30 days after sowing (DAS), 45 DAS, 60 DAS and 75 DAS. Data presented are means ± standard errors (*n* = 10, biological replicates). Different letters (a, b, c) indicate significant differences from control in all combinations (Tukey’s test, *P* ≤ 0.05).

|  |  | Chla | Chlb | CAR | TSS | MDA | PL | TPC | Prot | Urease | Leg Hb |
| --- | --- | --- | --- | --- | --- | --- | --- | --- | --- | --- | --- |
| **CN** | 30DAS | 4.98±0.29a | 1.03±0.06a | 1.02±0.06a | 9.2±0.55a | 0.63±0.03a | 2.83±0.14a | 1.65±0.08a | 1.36±0.06a | 7.22±0.36a | 1.26±0.29a |
| | 45DAS | 6.21±0.37a | 1.38±0.08a | 1.96±0.11a | 12.51±0.75 | 1.23±0.06a | 3.56±0.17a | 2.56±0.12a | 1.59±0.07a | 8.60±0.43a | 2.54±0.08a |
| | 60DAS | 6.77±0.40a | 1.40±0.08a | 2.03±0.12a | 20.02±1.20 | 1.54±0.07a | 4.65±0.23a | 3.66±0.18a | 2.34±0.11a | 7.02±0.35a | 4.93±0.07 |
| | 75DAS | 6.42±0.38a | 3.41±0.20a | 1.75±0.10a | 20.36±1.22 | 1.88±0.09a | 6.56±0.32a | 4.56±0.22a | 3.20±0.16a | 4.02±0.20a | 6.09±0.19 |
| **Cd** | 30DAS | 2.25±0.13b | 2.63±0.15b | 1.62±0.09b | 5.2±0.31 | 0.85±0.04b | 8.56±0.42b | 1.99±0.09a | 3.58±0.17b | 6.74±0.33a | 0.79±0.03 |
| | 45DAS | 3.55±0.21b | 2.28±0.13b | 2.02±0.12a | 7.26±0.43 | 1.65±0.08b | 10.56±0.52b | 2.56±0.12a | 2.15±0.10b | 5.47±0.27b | 2.69±0.17 |
| | 60DAS | 3.42±0.20b | 4.58±0.27b | 2.59±0.15b | 9.42±0.56 | 2.88±0.14b | 17.23±0.86b | 4.56±0.22b | 3.22±0.16b | 4.91±0.24b | 2.14±0.04 |
| | 75DAS | 3.81±0.22b | 3.58±0.21a | 1.75±0.10a | 11.96±0.71 | 3.01±0.15b | 19.36±0.96b | 5.66±0.28a | 3.06±0.15a | 4.52±0.22a | 2.77±0.29 |
| **GA3** | 30DAS | 5.90±0.35a | 2.67±0.16b | 3.39±0.20 | 8.7±0.52c | 0.84±0.04b | 4.56±0.22c | 2.15±0.10b | 1.78±0.08a | 7.97±0.39a | 1.02±0.04 |
| | 45DAS | 6.44±0.38a | 3.30±0.19b | 2.49±0.14b | 11.55±0.69 | 1.65±0.08b | 5.60±0.28c | 2.66±0.13a | 2.19±0.10b | 7.52±0.37a | 4.02±0.29 |
| | 60DAS | 7.25±0.43a | 2.62±0.15c | 2.37±0.14a | 26.6±1.59 | 2.00±0.10a | 12.56±0.62c | 3.36±0.16a | 3.63±0.18b | 7.66±0.38a | 5.42±0.15 |
| | 75DAS | 7.72±0.46c | 5.73±0.34b | 1.88±0.11b | 31.24±1.87 | 2.15±0.10a | 16.23±0.81b | 4.56±0.22a | 3.56±0.17a | 7.85±0.39b | 4.58±0.16 |
| **Cd+GA3** | 30DAS | 3.94±0.23c | 4.00±0.24c | 1.63±0.09 | 5.6±0.33b | 0.72±0.03a | 15.23±0.76d | 1.89±0.09b | 1.42±0.07a | 10.01±0.50b | 0.91±0.03 |
| | 45DAS | 3.97±0.23b | 3.41±0.20c | 2.84±0.17b | 8.50±0.51 | 1.35±0.06a | 19.65±0.98d | 2.44±0.12a | 2.45±0.12b | 9.54±0.47a | 2.68±0.08 |
| | 60DAS | 4.25±0.25b | 3.51±0.21c | 3.17±0.19c | 18.93±1.13 | 2.14±0.10c | 20.12±1.00b | 3.66±0.18a | 3.77±0.18c | 9.35±0.46c | 6.04±0.58 |
| | 75DAS | 4.26±0.25d | 4.09±0.24c | 2.90±0.17b | 21.67±1.3 | 2.04±0.10a | 21.36±1.06b | 4.33±0.21a | 2.70±0.13b | 5.12±0.25c | 8.16±0.42 |

### 3.4. GA3 improves plant biomass under Cd stress

Plant biomass is a good indicator of plant fitness under abiotic stress. In present study, Cd stressed plants showed 49% reduction in fresh leaf biomass (FLB) and 29% in dry leaf biomass (DLB) compared to control **(Figure 3a-b)**. GA3 could improve FLB and DLB by 24% and 63% in Cd stressed plants over Cd treated plants alone **(Figure 3a-b)**. GA3 alone improved FLB and DLB non-significantly compared to control plants. Stem fresh (SFB) and dry biomass (SDB) showed reduction by 17% and 48% in Cd stressed plants over control. GA3 supplementation to Cd stressed plants witnessed increased SFB (13%) and SDB (9.5%) compared to Cd alone treated plants **(Figure 3c-d)**. GA3 alone could improve SFB (35%) and SDB (55%) over untreated control plants. Root biomass is a key determinant for successful survival of a plant under abiotic stresses **(Figure 3e-f)**. Under Cd stress, 18% and 25% reduction in fresh root biomass (FRB) and dry RB was noted over control. GA3 could improve both FRB and DRB by 30% and 23% in Cd stressed plants over Cd treated plants. GA3 alone improved FRB and DRB by 30% and 23% over control plants **(Figure 3e-f)**.

**Figure 3.**
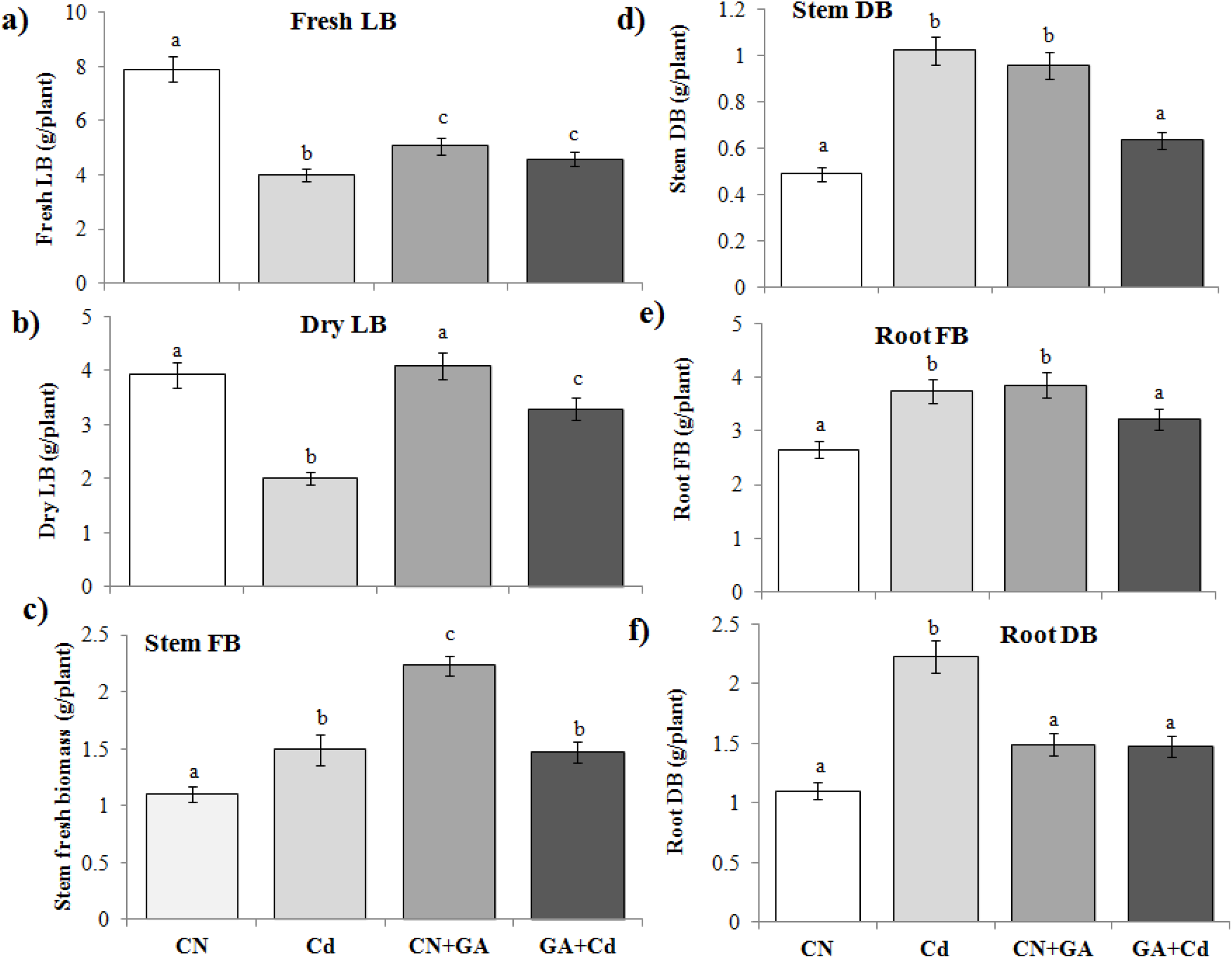
Gibberellins (GA3) enhance biomass of mung bean plants under cadmium (Cd) stress in soil. Effect of GA3 with or without cadmium (Cd) stress on fresh leaf biomass (FLB) (a), dry leaf biomass (DLB) (b), stem fresh biomass (c), stem dry biomass (d), root fresh biomass (e) and root dry biomass (f) of mung bean plants grown in soil in pots at time of harvest. Control (CN), cadmium (Cd) stress, Cd+ gibberellins (GA3) and GA3 alone conditions. Data presented are means ± standard errors (*n* = 10, biological replicates). Different letters (a, b, c) indicate significant differences from control in all combinations (Tukey’s test, *P* ≤ 0.05).

### 3.5. GA3 improves photosynthetic pigments and soluble sugar under Cd stress

Photosynthetic pigments suffered damage under Cd stress such that a consistent decrease in Chl *a* content at all the observation points (30, 45, 60 and 75 DAS), with maximum reduction (55%) occurred at 30 DAS compared to control **(Table 1)**. GA3 could improve Chl *a* content consistently, with maximum increase (75%) recorded at 30 DAS over Cd alone treated plants. GA3 alone improved Chl *a* content at all observation points, with maximum enhancement (20%) noted at 75 DAS compared to control plants **(Table 1)**. Contrary to Chl *a*, under Cd stress, Chl *b* content showed consistent increase at all observation points, with maximum increase 225% recorded at 60 DAS compared to control plants. Furthermore, GA3 could increase Chl *b* content throughout the experiment, with maximum improvement (52%) noted at 30 DAS over Cd alone treated plants **(Table 1)**. GA3 alone could improve Chl *b* content at all the observation points, with maximum increase (157%) being recorded at 30 DAS compared to control plants. For carotenoids (CARs), under Cd stress, content showed upward trend, with maximum increase (59%) noted at 30 DAS over control plants. GA3 improved CAR content maximally (66%) at 75 DAS compared to only Cd treated plants. GA3 alone could improve CAR content at all the observation points, with maximum increase (232%) recorded at 30 DAS compared to control plants **(Table 1)**. Total soluble sugars (TSS) amount was significantly reduced at 30, 45, 60 and 75 DAS in Cd stressed plants compared to control. GA3 supplied to Cd stressed plants significantly enhanced TSS by 81% and 100% at 60 and 75 DAS compared to Cd alone treated plants. GA3 alone could improve TSS significantly at 75 DAS over control plants **(Table 1)**.

### 3.6. GA3 lowers stress indices and boosts antioxidant system to combat Cd phytotoxicity in pot experiments

Stress indices such as malondialedhye (MDA, membrane damage indicator) and proline (PL) play significant role in evaluating the physiological status of a plant under abiotic stress. In present study, Cd stress was shown to increase MDA content at all the observation points, with most significant membrane damage (87%) recorded at 60 DAS compared to control plants **(Table 1)**. GA3 could bring decline in MDA content in Cd stressed plants maximally at 75 DAS over Cd treated plants. GA3 alone significantly increased MDA (34%) at 45 DAS compared to control plants. Under Cd stress, PL content rise sharply, with maximum increase (270%) at 60 DAS compared to control plants. GA3 supplementation to Cd stressed plants further improved PL content at all the observation points, with maximum increase (86%) recorded at 45 DAS over Cd alone treated plants. A consistent increase in PL occurred under GA alone, with maximum increase (170%) noted at 60 DAS over control plants **(Table 1)**.

Among antioxidants, total phenol content (TPC) increased maximally (25%) at 60 DAS in Cd stressed plants compared to control plants **(Table 1)**. GA3 applied to Cd stressed plants showed decline in TPC content at all the observation points, with maximum reduction (24%) noted at 75 DAS over Cd alone treated plants. In comparison, GA3 alone treated plants showed maximum increase in TPC (30%) at 30 DAS **(Table 1)**. Cd stressed plants showed significant increase in total GSH content (640%) compared to control plants (*p >* 0.05). GA3 application to Cd stressed plants lowered total GSH content by 26% over only Cd treated plants. GA3 alone could improve total GSH by 1000% compared to control plants **(Table 1).**

### 3.7 GA3 reduces accumulation, translocation and localization of Cd ions

Compared to control plants, Cd stressed plants showed 61%, 126% and 88% higher accumulation of Cd in leaf, shoot and root tissue (**Fig. 4a**). GA3 application to Cd stressed plants reduced accumulation of Cd by 35%, 34% and 6% in leaf, shoot and root tissues compared to Cd alone treated plants (**Fig. 4a**). About 75% and 65% accumulation of Cd was noted in pods and seeds of Cd stressed plants compared to control plants. GA3 application reduced Cd accumulation by 67% and 89% in pods and seeds when compared to Cd alone treated plants. Bioaccumulation factor (BAF) from soil to root transfer of Cd showed significant increase 14.48-fold over soil concentration of Cd alone (**Fig. 4b**). GA3 application to Cd stressed plants showed significant reduction of BAF of roots from soil by 7.79-fold compared to BAF in Cd alone (14.48-fold) (**Fig. 4b**). Translocation factor (TF) from shoot-leaf showed no significant change in Cd alone and Cd stressed plants supplemented with GA3 (**Fig. 4c**). For histological localization of Cd^2+^ ions, white crystalline precipitates of Cd oxalate were observed in the transverse sections (T.S.) of the stem of plants under Cd stress. Localization was more pronounced in the cortex areas of Cd stressed plants, while Cd oxalate crystals were very few in GA3 applied Cd plants **(Fig. 4d-g)**.

**Figure 4.**
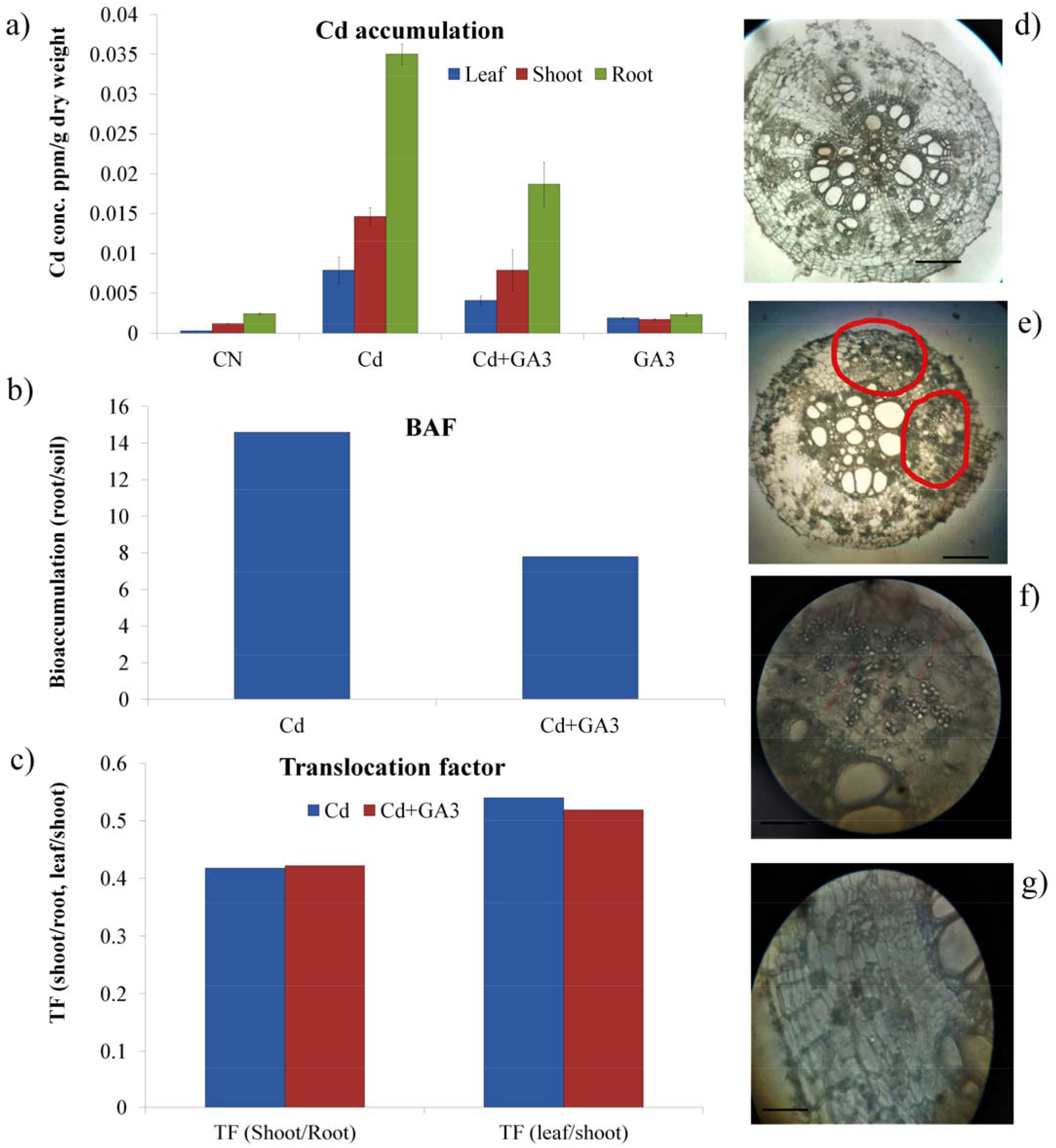
Gibberellins (GA3) reduce accumulation, translocation and localization of cadmium (Cd) in mung bean plants. Effect of GA3 on cadmium (Cd) accumulation (ppm/g dry weight) in leaf, shoot and root tissue (a), bio accumulation factor (BAF, Cd accumulation root/soil) (b), translocation factor (shoot/root and leaf/shoot) (c) and histological localization of Cd ions as Cd-oxalate white crystals in stem tissue transverse sections, Cd alone (d-e) and GA3 pus Cd (f-g) of mung bean plants grown in soil in pots at time of harvest. Control (CN), cadmium (Cd) stress, Cd+ gibberellins (GA3) and GA3 alone conditions. Data presented are means ± standard errors (*n* = 10, biological replicates) for a-c.

### 3.8. GA3 modulate genes regulating Cd uptake and transport

Expression of genes inducing Cd^2+^ ions sequestration was enhanced under Cd stress compared control. Such that 2-fold increase in expression of *VrPCS1* and *VrPCS2* was noted in Cd stressed seedlings compared to control. GA3 application further elevated expressions of *VrPCS1* and *VrPCS2* over Cd treatment (**Fig. 5a-b**). GA3 alone could not modulate expressions of these genes significantly compared to control. About 10-fold increase in uptake of Cd ions under Cd stressed plants over control was linked with 3-fold increase in expression of *VrIRT1* over control plants. GA3 application lowered Cd accumulation in young plants by 4-fold, this reduction in Cd ions accumulation could be attributed to 2-fold decline in expression of *VrIRT1* compared to only Cd treated seedlings (**Fig. 5c**). GA3 alone also improved *VrIRT1* expression over control though insignificantly when compared to control. Elevated expression of *VrCDR* noted Cd stress alone compared to control. GA3 application to Cd stress further improved *VrCDR* expression when compared to only Cd treatment **(Figure 5d)**.

**Figure 5.**
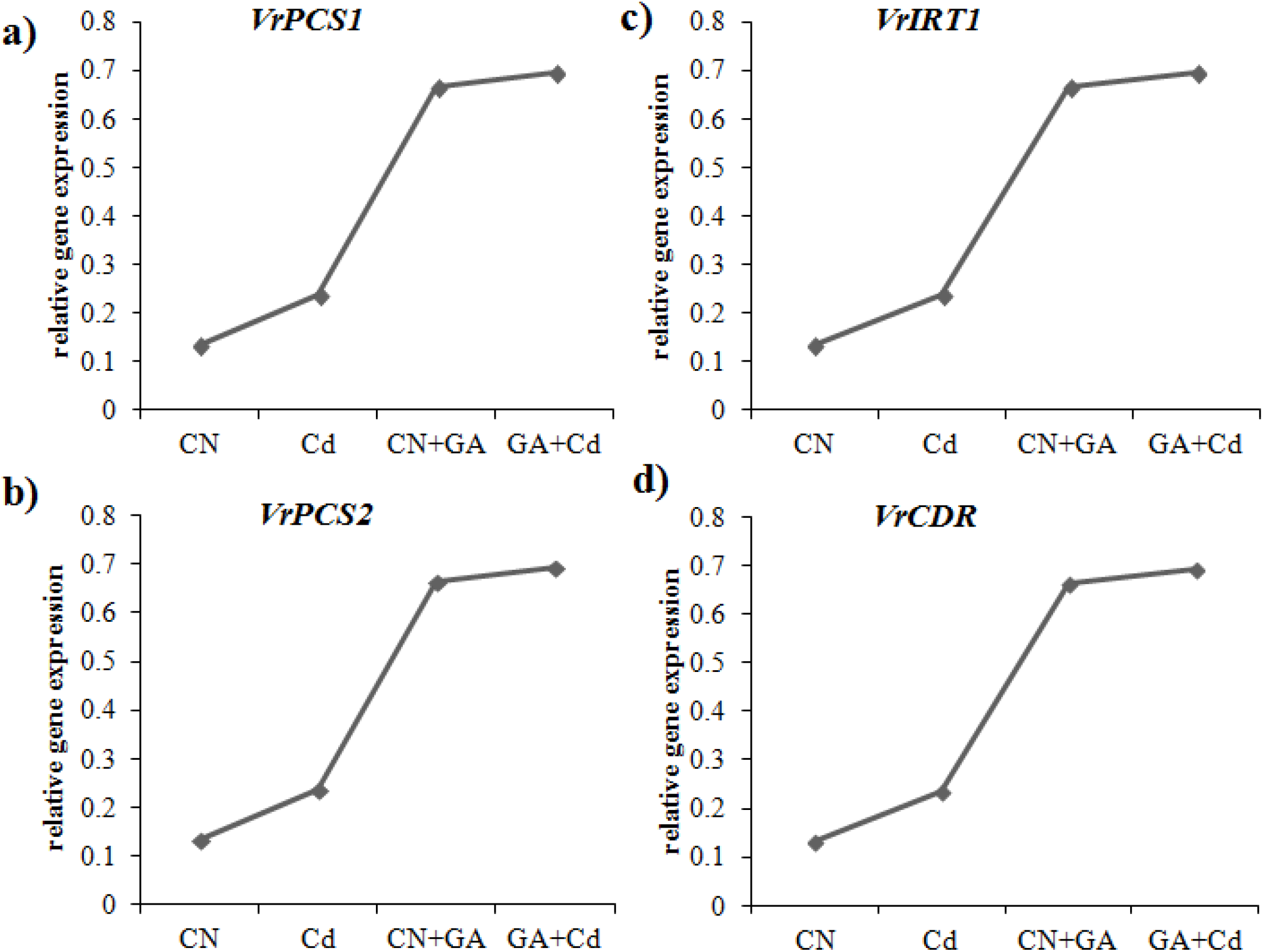
(a-d) Gibberellins (GA3) modify expression profile of heavy metal stress tolerance genes in mung bean plants. Effect of GA3 on the relative gene expression of *Vigna radiata phytochelatin synthase 1* (*VrPCS1*), *VrPCS2*, *VrIron transporter* 1 (*VrIRT1*) and *Vr cadmium response* (*VrCDR*) in young plants subjected to Control (CN), cadmium (Cd) stress, Cd+ gibberellins (GA3) and GA3 alone conditions in paper roll method. Data presented are means ± standard errors (*n* = 3, biological replicates).

### 3.9. GA3 improves agronomic traits affected by Cd stress

A visible reduction in pod length and seed size was observed under Cd stress, while GA3 application could mitigate negative impacts of Cd stress on pod length and seed size (**Fig. 6**). For pod length, Cd treated plants showed 21% reduction, while GA3 improved pod length by 11% in Cd stressed plants over Cd alone **(Figure 7a)**. Cd treated plants showed 8% reduction in pod diameter compared to control plants. GA3 could improve pod diameter by 17% in Cd stressed plants **(Figure 7b)**. GA3 applied alone increased pod diameter by 58% over control plants **(Figure 7b)**. Approximately 34% reduction in pod number/plant was recorded in Cd stressed plants over control plants. About 25% increase in pod number/plant was recorded for GA3 applied Cd stressed plants over Cd alone treated plants. No significant change in pod number/plant was recorded for alone GA3 treated plants over control plants **(Figure 7c)**. About 49% and 52% reduction in pod fresh and pod dry weight was noted in Cd stressed plants compared to control plants. GA3 in Cd stressed plants showed significant improvement in pod fresh (57%) and pod dry weight (107%) over Cd alone treated plants **(Figure 7d-e)**. Interestingly, 108% and 237% increase in pod fresh and pod dry weight was recorded for alone GA treated plants compared to control plants **(Figure 7d-e)**.

**Figure 6.**
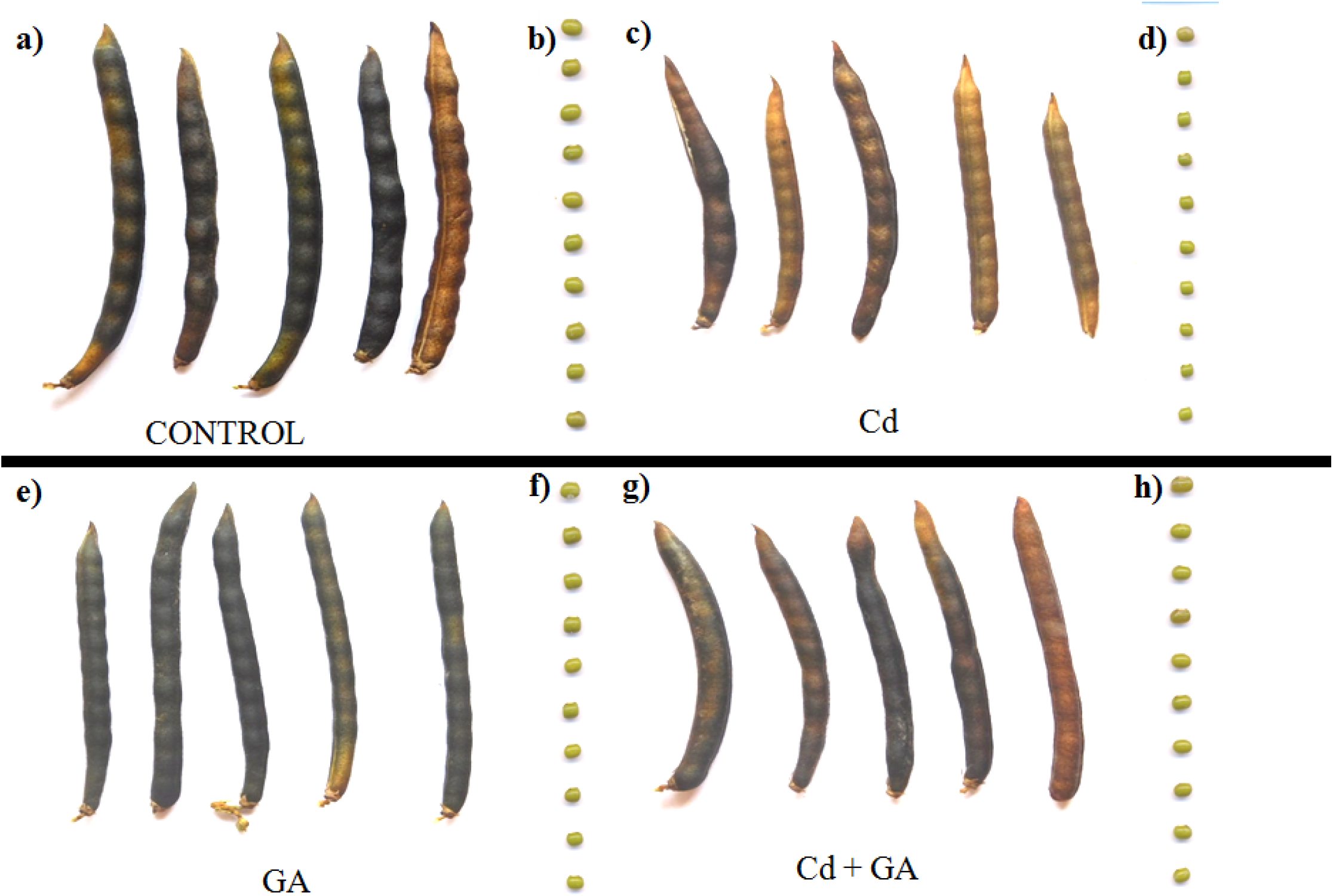
(a-h). Gibberellins (GA3) improve pod and seed size (illustration images) under Cd stress. Control (CN, a), cadmium (Cd, b) stress, GA3 (c) and Cd+ and GA3 conditions.

**Figure 7.**
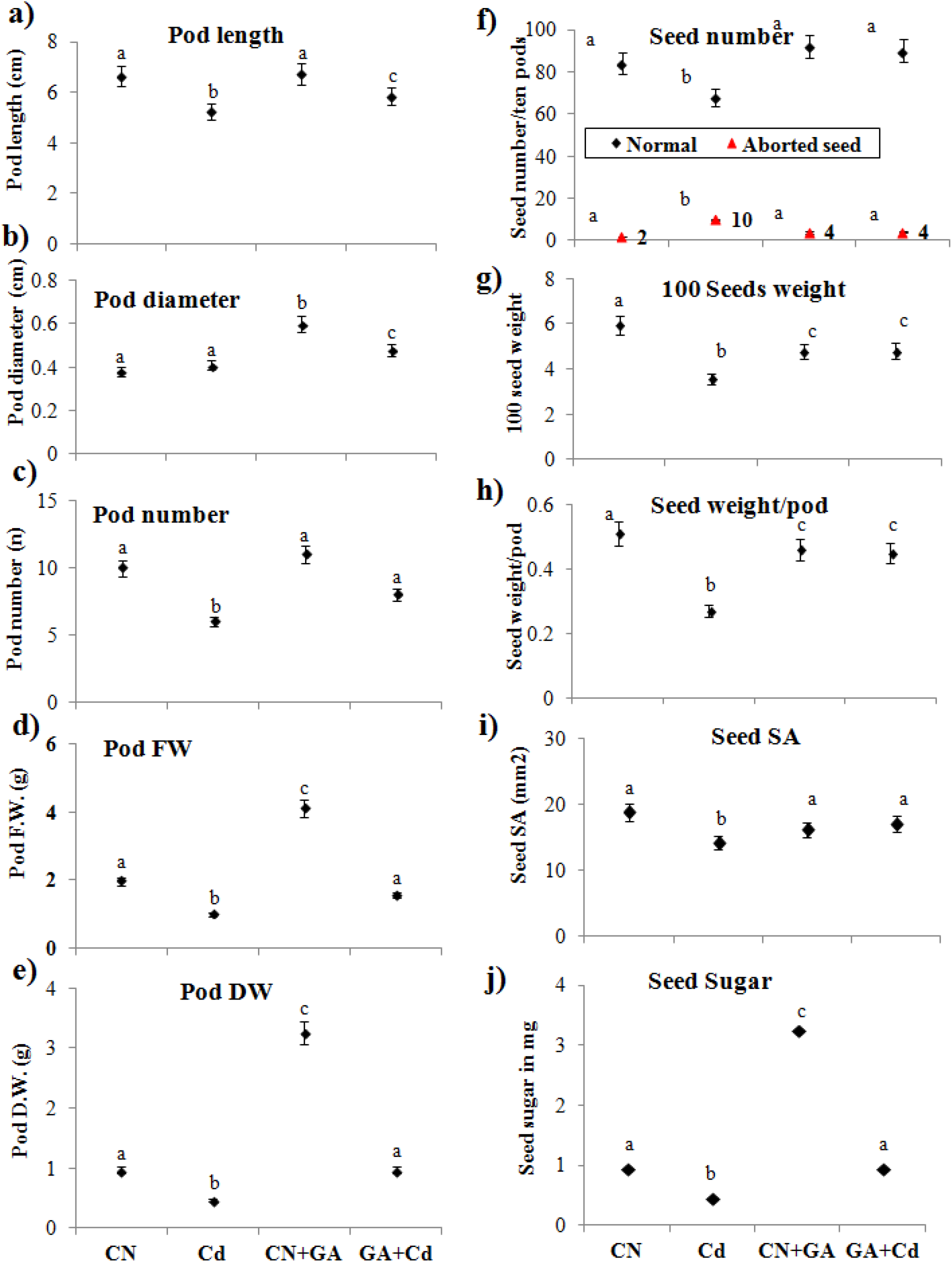
Gibberellins (GA3) improves agronomic attributes of mung bean plants under cadmium (Cd) stress in soil. Effect of GA3 with or without cadmium (Cd) stress on pod length (a), pod diameter (b), pod number/plant (c), pod fresh weight/plant (Pod FW, d), pod dry weight (Pod DW, e), seed number/ten plants (f), hundred seeds weight (g), seed weight/pod (h), seed surface area (i) and seed sugar (j) of mung bean plants grown in soil in pots at time of harvest. Control (CN), cadmium (Cd) stress, Cd+ gibberellins (GA3) and GA3 alone conditions. Data presented are means ± standard errors (*n* = 10, biological replicates). Different letters (a, b, c) indicate significant differences from control in all combinations (Tukey’s test, *P* ≤ 0.05).

For seed, Cd stress was able to influence several important seed attributes viz. seed number/ten pods, 100 seeds weight, seeds weight/pod, seed width (W), seed length (L), seed thickness (T), seed surface area (SA), mean geometric diameter (D_g_), sphericity (Φ%), seed volume (V) and seed germination percentage (%G). Such that 19%, 40% and 47% reduction in seed number/ten pods, 100 seeds weight and seed weight/pod was recorded in Cd stressed plants over control plants. However, GA3 application in Cd stressed plants showed 32%, 35% and 67% improvement in these seed attributes over alone Cd treated plants **(Figure 7f-h)**. GA3 alone improved seed number/ten pods by 9.5% and reduced 100 seeds weight and seed weight/pod by 20% and 10% respectively compared to control plants **(Figure 7f-h)**. Seed surface area ()For seed geometric attributes, about 13%, 15%, 14% and 16% reduction in W, L, T and SA was recorded in Cd stressed plants compared to control plants **(Figure 7i, Table 2)**. GA3 application to Cd stressed plants improved W, L, T and SA by 8.47%, 15.29%, 8.23% and 13.27% respectively over Cd alone treated plants. Similarly, 13.77% and 35.89% reduction in D_g_, and V was recorded in Cd stressed plants over control plants **(Table 2)**. GA3 application in Cd stressed plants showed 10.46% and 34.79% improvement in D_g_, and V over Cd alone treated plants. Contrary, GA3 application alone reduced D_g_ by 8.31% and V by 22.91% compared to control plants alone. While, no significant change in Φ insignificant decrease of 4% was recorded for GA3 applied Cd stressed plants compared to alone Cd treated plants **(Table 2)**. Seed protein content under Cd stress declined 36% compared to control seeds. Application of GA3 in Cd stressed plants further declined seed protein content by 20% over Cd treated plants. Interestingly, seed protein content declined by 35% in GA3 alone treated plants over control plants **(Table 2)**. Seed TPC in Cd stressed plants showed significant increase (241%) over control plants. GA3 applied Cd stressed plants showed 92% reduction in seed TPC over TPC of seeds of Cd alone treated plants. About 43.46% reduction in seed TPC was noted in alone GA treated plants compared to control **(Table 2)**.

**Table 2.**
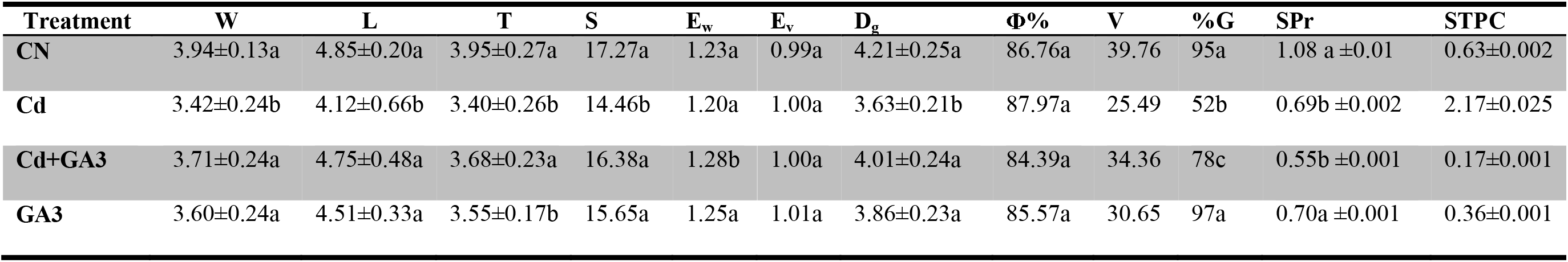
Effects of cadmium (Cd) stress with or without gibberellins (GA3) on seed geometric attributes viz. width (W), length (L), thickness (T), Sphericity (S), Ew, Ev, geometric mean (Dg), seed shape (Φ%), volume of the grain (V), gravity (%G), SPr, STPC of seeds of mung bean plants subjected to control (CN), cadmium stress (Cd), gibberellins (GA3), Cd+GA3 at 30 days after sowing (DAS), 45 DAS, 60 DAS and 75 DAS. Data presented are means ± standard errors (*n* = 10, biological replicates). Different letters (a, b, c) indicate significant differences from control in all combinations (Tukey’s test, *P* ≤ 0.05).

| Treatment | W | L | T | S | E<sub>w</sub> | E<sub>v</sub> | D<sub>g</sub> | $\Phi\%$ | V | %G | SPr | STPC |
| --- | --- | --- | --- | --- | --- | --- | --- | --- | --- | --- | --- | --- |
| CN | 3.94±0.13a | 4.85±0.20a | 3.95±0.27a | 17.27a | 1.23a | 0.99a | 4.21±0.25a | 86.76a | 39.76 | 95a | 1.08 a ±0.01 | 0.63±0.002 |
| Cd | 3.42±0.24b | 4.12±0.66b | 3.40±0.26b | 14.46b | 1.20a | 1.00a | 3.63±0.21b | 87.97a | 25.49 | 52b | 0.69b ±0.002 | 2.17±0.025 |
| Cd+GA3 | 3.71±0.24a | 4.75±0.48a | 3.68±0.23a | 16.38a | 1.28b | 1.00a | 4.01±0.24a | 84.39a | 34.36 | 78c | 0.55b ±0.001 | 0.17±0.001 |
| GA3 | 3.60±0.24a | 4.51±0.33a | 3.55±0.17b | 15.65a | 1.25a | 1.01a | 3.86±0.23a | 85.57a | 30.65 | 97a | 0.70a ±0.001 | 0.36±0.001 |

### 3.10 Pearson correlation analysis

A significant linear correlation (*r*) for stem height, stem diameter and leaf number was noted for Cd stress alone **(Table 3)**. Pearson’s correlation coefficient (*r*) for shoot traits under Cd stress showed linear correlation; however significant positive correlation was noted for stem diameter **(Table 3)**. Negative correlation was noted for root length under Cd stress **(Table 3)**. GA3 applied to Cd stressed plants showed positive correlation for stem diameter, leaf number and root length, while a negative correlation noted for stem height and leaf surface area **(Table 3)**. For agronomic traits, seed number, seed weight, seed weight/pod showed positive correlation with Cd stress, while a negative correlation was found for seed surface area and total GSH **(Table 3)**. While all agronomic traits except seed number showed negative correlation between GA3 applied Cd stressed plants and Cd alone treated plants **(Table 3)**.

**Table 3.** Pearson correlation coefficient (*r*) for ten traits of plant fitness (stem height, stem diameter, leaf number, leaf surface area and root length) and of agronomical attributes (seed number, seed weight, seed weight/pod, seed surface area, total GSH) for Control versus Cd, Control versus Cd+GA3 and Control versus GA3. Number of biological replicates (*n =* 13) across treatments except total GSH, where *n* = 10. Asterisk * represents Correlation is significant at the 0.05 level (2-tailed).

| **Pearson correlation for morphometric traits** |  |  |  |  |  |
| --- | --- | --- | --- | --- | --- |
|  | *Stem height* | *Stem diameter* | *Leaf number* | *Leaf surface area* | *Root length* |
| **CN** | 1 | 1 | 1 | 1 | 1 |
| **Cd** | 0.364 | 0.593* | 0.014 | 0.447 | -0.289 |
| **Cd+GA3** | -0.050 | 0.875 | 0.582* | -0.061 | 0.473 |
| **Pearson correlation for agronomic traits** |  |  |  |  |  |
|  | *Seed number* | *Seed weight* | *Seed weight/pod* | *Seed surface area* | *Total GSH* |
| **CN** | 1 | 1 | 1 | 1 | 1 |
| **Cd** | 0.127 | 0.075 | 0.295 | -0.156 | -0.415 |
| **Cd+GA3** | 0.088 | -0.413 | -0.400 | -0.032 | -0.849 |

## Discussion

Cadmium (Cd) is a non essential heavy metal known to affect human health and plant growth and metabolism (Hu *et al*., 2020; Sanaei *et al*., 2020). Being highly active Cd, its excess in soil often accounts for perturbed metabolism and growth defects leading to reduced yield in crops of commercial importance (Shanying *et al*., 2017). Decrease in shoot and root lengths and fresh and dry biomass due to heavy-metal stress has been reported in mungbean (Ghani, 2010), *Arachis hypogea* (Lu *et al*., 2013), and sunflower (Mohammadzadeh *et al*., 2017). Decrease in plant height under Cd stress in current study (Fig. 1) can be supported with similar observations recorded in *Vigna radiata* (L.) Wilzeck cultivars PDM-139 and K-851 subjected to 0.2 mg/L of Cd exposure showed significant reduction in plant height (Kumari *et al*., 2011). However, improved plant height recorded in Cd stressed plants upon application of GA3 compared to Cd stressed plants could be attributed to positive impact of GA3 on inducing plant growth (Fig. 1). Our observations are supported by similar findings, wherein application of GA (50 µM) to *Brassica napus* L. grown in Cd (0, 50 and 100 μM) hydroponics could increase plant height compared to only Cd hydroponics (Meng *et al*., 2009). Similar to our observations of reduced leaf number and leaf SA in mung bean plants under Cd stress, reduction in leaf number and leaf SA have been reported in *V. radiata* (L.) Wilczek subjected to increased levels of Cd in soil medium (Ghani and Wahid, 2007). Improved leaf SA of Cd stressed plants applied with GA3 (Fig. 1) could be linked with established impact of GAs on the leaf elongation rate (LER) *via* -expansins, one β xyloglucan endotransglycosylase (Xu *et al*., 2016). Reductions in both fresh and dry biomasses of Cd stressed plants in current study could be attributed to Cd interference in normal functioning of enzymes of Calvin cycle, CO2 fixation, phosphorus and carbohydrate metabolism leading to stunted plant growth and inhibition of photosynthesis (Gill and Narendra, 2011; Tiwari and Lata, 2018).

Plants often respond to soil HM stress by reducing root length and root surface area, so as to minimize the root area exposition to HM pollution (Rizvi & Khan, 2017). Current study also noted decrease in root length and root surface area and root biomass both in young plants (Experiment 1) and mature plants (Experiment 2) (Fig. 1, 2 and 3). Role of Cd in decreasing nodulation and activity of nitrogen metabolizing enzymes has been established (Alyemeni *et al*., 2016). Higher concentrations of Cd have been observed to induce nodule senescence and decrease in legHb content in leguminous plants (Alyemeni *et al*., 2016). Contrary to earlier reports, Cd stress was observed to increase nodule number/plant over control plants, interestingly this increase in nodule number/plant was found to be associated with significant reduction in nodule diameter and nodule surface area (Fig.1). However, application of GA3 was able to improve nodule diameter at all observation points with maximum increase recorded at 75 DAS compared to only Cd stressed plants (Fig.1). Similar to observations reported earlier, Cd stress was able to lower the nodule legHb content significantly over control plants; however, supplementation of GA3 to Cd stressed plants showed improvement in nodule legHb content (Table 2). This reduction in legHb content under Cd stress could be attributed to production of higher number of reactive oxygen species with detrimental impact on nodules (Balestrasse *et al*., 2004).

A positive correlation existed between accumulation of Cd in shoot tissue and its localization (Fig. 4a). Significantly accumulated Cd in Cd stressed plants followed shoot > root > leaf, while GA3 application could reduce Cd accumulation and root-shoot and shoot to leaf translocation significantly (Fig. 4 b). Similarly, accumulation of Cd was higher in pods than seeds, while GA3 application reduced this accumulation by a significant factor (Fig. 4b). This significant reduction in Cd accumulation in GA3 treated plant tissues and its translocations from root to shoot and shoot to leaf could be linked with elevated expressions of genes regulating Cd uptake (*VgCDR1*), phytochelatin synthesis (*VgPCS1* and *VgPCS2*) and *VgIRT1* compared to Cd stressed plants (Fig. 5).

Reductions in photosynthetic pigments (Chl *a*, Chl *b* and CAR) in Cd stressed plants (Table 1) may be linked to inhibitory impact of Cd induced generation of ROS and lipid peroxidation and inhibiting Chl biosynthesis by competing with Mg ions consequently leading to damage of PS II (Rai *et al*., 2016). Application of GA3 ameliorated negative impact of Cd on photosynthetic pigments *via* reducing the production of ROS and damage to cell membrane by improving antioxidant activity of Cd stressed plants compared to Cd alone treated plants (Table 1). Exogenous applications of GA3 have been shown to improve contents of photosynthetic pigments by stabilizing and inhibiting the dissociation of the light-harvesting complex from PS II core, under Cu stress in *Helianthus annuus* L. (Ouzounidou and Ilias, 2005).

In current study, Cd stressed plants showed increase in membrane damage as indicated by enhanced levels of MDA throughout the plant’s life cycle after imposition of Cd stress (Table 1). Application of GA3 was observed to reduce membrane damage in Cd stressed plants probably via reducing the production of ROS compared to Cd stressed plants (Table 1). Findings are in consonance with Cd induced membrane damage and higher production of superoxide radicals and membrane lipid peroxidation products recorded in seedlings of *Vigna aconitifolia* L. (Vijendra *et al*., 2016). Enhanced levels of PL in Cd stressed plants could be linked with effective HM stress management (Table 1); moreover, GA3 induced further increase in PL content in Cd stressed plants can be associated with reducing ROS production. GA3 induced increase in PL production and its implication in bud outbreak has been recorded in *Prunus avium* (sweet cherry) (Cai *et al*., 2019).

Phenolic acids play pivotal roles in abiotic stress management, particularly by neutralizing or scavenging ROS. TPC is a good indicator of total phenolics content in a plant system under abiotic stresses in plants. In current study, enhanced TPC in Cd stressed plants could be linked with plant’s inbuilt response towards enhanced ROS production and their management (Table 1). While GA3 induced small decrease in TPC of Cd stressed plants could be attributed towards reduction of ROS levels and returning of normal growth conditions compared to Cd stressed plants (Table 1). Enhanced levels of total phenolics in GA treated *Vitis vinifera L*. cv. Muscat have been recorded to play important roles in development of grapevine tissues (Tian *et al*., 2011). Soil HM pollution has been shown to affect activity of nitrogen metabolism enzymes such as urease and nitrification dynamics in the soil-rhizosphere zone (Yan *et al*., 2013). At low concentrations (0.5 mg/kg soil), Cd^2+^ cations were able to up-regulate the activity of urease enzyme; though at higher concentration activity got declined considerably (Yan *et al*., 2013). Similar observations were also recorded for Cd stressed plants and elevated activity of urease in GA3 applied plants (Table 1).

Cd stress has been observed to reduce agronomic traits of *V. radiata* L (Ghani, 2010), such that plants subjected to 3, 6, 9 and 12 mg/kg soil Cd toxicity showed reduction in the number of pod/plant, pod length, pod fresh weight, pod dry weight, number of seeds/pod and number of seeds/plant (Ghani, 2010). Current study also witnessed reduction in pod size, pod length, pod diameter, pod fresh and dry weights, seed number/pod and plant, seed weight under Cd stress (Fig. 6 and 7). In addition, reduced seed surface area, sugar content, protein content and geometric attributes were also noted under Cd stress (Fig. 7 and Table 2). GA3 supplementation was able to significantly mitigate negative impact of Cd stress on all the pre mentioned agronomic attributes (Fig. 7 and Table 2).

Pearson correlation analysis also revealed negative impact of Cd stress on agronomic and growth parameters of vigna plants (Table 3). Moreover, GA3 application was seen to improve negative impact of Cd stress on growth and agronomic traits considerably. Findings thus propose, application of GA3 for significantly reducing the Cd stress associated damages in mung plants.

## Acknowledgements

Authors would like to thank Department of Botany, University of Jammu for extending laboratory facilities and technical support. The research work was financially supported by UGC-DRS-SAP-PHASE-II, New Delhi, India and CSIR-Junior Research Fellowship, Government of India to Urfan Mohammad. There are no conflicts of interest among the authors of this manuscript.

## Authors Contributions

S.P. conceived and designed the experiments; H.R.H., M.U., S.S. and D.K. performed experiments. N.S.Y. and S.P. performed data analysis, prepared figures and wrote the article.

